# Vocal communication is seasonal in social groups of wild, free-living house mice

**DOI:** 10.1101/2024.10.07.617064

**Authors:** N Jourjine, C Goedecker, B König, AK Lindholm

**Affiliations:** Department of Molecular & Cellular Biology, Department of Organismic & Evolutionary Biology, Center for Brain Science, Museum of Comparative Zoology, Harvard University, 16 Divinity Avenue, Cambridge, MA 02138, USA; Department of Evolutionary Biology & Environmental Studies, University of Zürich, Winterthurerstrasse, 190 8057 Zürich, Switzerland

## Abstract

House mice (*Mus musculus domesticus*) are among the most widely studied laboratory models of mammalian social behavior, yet we know relatively little about the ecology of their behaviors in natural environments. Here, we address this gap using radiotelemetry to track social interactions in a population of wild mice over 10 years, from 2013 to 2023, and interpret these interactions in the context of passive acoustic monitoring data collected from August 2022 until November 2023. Using deep convolutional neural networks, we identify 1.3 million individual vocalizations and align them in time with continuously collected telemetry data recording social interactions between individually identifiable mice. We find that vocalization is seasonal and correlated with long-term dynamics in features of social groups. In addition, we find that vocalization is closely associated in time with entrances to and exits from those groups, occurs most often in the presence of pups, and is correlated with how much time pairs of mice spend together. This work identifies seasonal patterns in the vocalizations of wild mice and lays a foundation to investigate the social role of acoustic communication in wild populations of a classic laboratory model organism.

---

Sociality is a defining feature of the human experience, but it is not unique to our species. Indeed, social behaviors are early hallmarks of vertebrate evolution^1^ and have diversified across distantly related taxa^2–5^, where they lead to the formation of population-wide social structures (i.e., “social networks”) whose features impact the survival and reproduction of their members^6,7^. How this social structure arises from dynamic interactions between individuals remains an open question in behavioral ecology.

Social interactions require communication among partners, and acoustic signaling is one such mode of communication that is pervasive among animals, where it serves diverse social roles ranging from the maintenance of territorial boundaries^8^ to group cohesion^9^ and many others^10^. It is therefore a useful context to understand how social interactions give rise to social structure in animal populations^11^. Doing this is challenging, however, because it requires monitoring social groups at spatial and temporal scales that are often large (e.g., the area occupied by a population over months or years) while recording acoustic signals taking place at scales that are necessarily small (e.g., the location of an individual as it vocalizes). Advances in hardware^12–14^ and software^15–18^ for passive animal tracking have made progress toward overcoming this challenge. Nonetheless, the unique ecologies of many social species make it infeasible to simultaneously track social groups and individual vocal behaviors at the population scale over ecologically meaningful time periods.

House mice (*Mus musculus domesticus*) may be one exception^19^. House mice are human-commensals that typically live at high population density in small areas near human activity, making it feasible to track nearly all individuals in a population in a single location^20^. In the wild and in semi-natural enclosures, they live in social groups^21–25^ and are also highly vocal^26,27^, producing two major call types that fall into the ultrasonic range (“USVs”) and human-audible range (“squeaks”)^28,29^. However, the social role of vocalization in this species has been primarily studied in a limited number of laboratory contexts such as mating^30,31^ and neonatal isolation^32^. While this work has uncovered fundamental neural mechanisms that support vocalization in these specific social contexts^33^, it is likely that they are a subset of the social contexts in which acoustic communication matters for survival and reproduction in natural habitats. As a result, the role that vocal communication plays in the social lives of wild mice remains unclear.

Here, we simultaneously record vocal behaviors and social interactions in a population of wild house mice living in a barn in the Swiss countryside. Using continuous tracking of individuals as they interact over the course of 10 years, we find that mice occupy seasonally dynamic social groups in which females play a central role, consistent with previous work examining shorter time scales^34^. Using passive acoustic monitoring of the same population over a 15-month period between August 2022 and November 2023, we further find that the amount of time mice spend vocalizing is seasonal, corresponds to events affecting the composition of social groups, and is correlated with how much time pairs of mice spend together. Taken together, these findings identify potential social roles for acoustic communication in seasonally dynamic social networks of wild mice.

## Results

We focused on a population of house mice (*Mus musculus domesticus*) inhabiting a barn in a wooded agricultural landscape about 20 km from Zürich, Switzerland (Figures 1A). The barn contains several small (15cm x 15cm), cylindrical boxes each fitted with a single, 4.4 cm diameter ground-level entryway and two radio-frequency identification (RFID) readers (“RFID boxes”, Figures 1B, C), in additional to a set of low internal walls and an entry area containing a desk and computer (Figure 1D: there were 40 RFID boxes until February 2021, when the number was intentionally reduced by half). It also provides some protection from predators and contains *ad libitum* food and water (see methods for details), making it an ideal habitat for house mice that recapitulates the human-made infrastructure they typically occupy, while also facilitating long-term monitoring of their behaviors in natural environments (Figure 1E).

**Figure 1.**
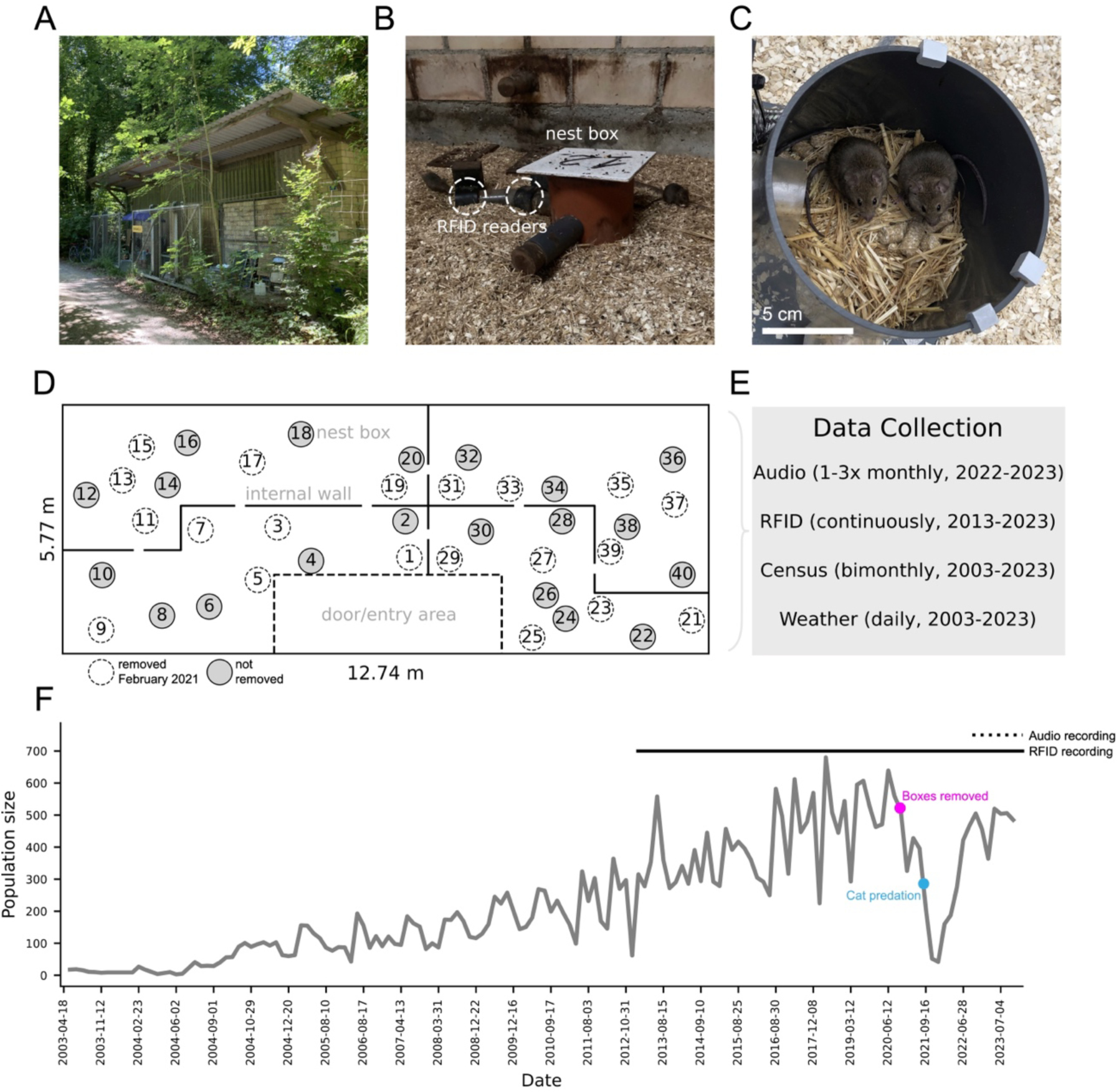
Long term monitoring of a wild house mouse population. **(A)** The outside of the study site, a barn located at the edge of a forest in an agricultural landscape **(B)** An RFID box inside the barn equipped with an entrance tunnel and RFID readers. Two mice are shown, one entering the box and one sitting next to it **(C)** Two mice inside an RFID box with the lid removed. Scale bar: 5cm. **(D)** A schematic of the barn interior, indicating RFID box locations **(E)** Data collected from the barn and its mice from April 2003 to December 2023; Audio: continuous, 48 hour long audio recordings from individual RFID boxes; RFID: radio frequency identification readings from transpondered mice during each unique entrance and exit to any RFID box; Census: whole population census in which each mouse in the population is caught; Weather: daily meteorological statistics for the barn location **(F)** Size of the mouse population in the barn from 2003 to 2023. The large population crash in 2021 corresponds to a series of unintentional cat predation events as well as an intentional reduction in the number of RFID boxes from 40 (2003-2021) to 20 (2021-present, as shown in D).

Twelve mice from neighboring farms were introduced to the barn in 2002. These founders have since given rise to a growing resident population of approximately 400-600 adults, which experienced one major crash caused by a series of cat predation events in the Autumn of 2021, following the removal of half of the RFID boxes in February of 2021 (Figure 1D,F). Access to the barn is limited only by body size and mice are free to enter and exit. House mice are nonetheless the barn’s only mammalian inhabitant, as confirmed at censuses that have occurred bimonthly for the last 22 years. During these censuses, all individuals are caught, counted, and adults fitted a subcutaneous microRFID transponder (Trovan, Ltd., United Kingdom) if they do not already have one before being released back into the barn, where the timestamps of each individual’s entrances and exits from RFID boxes are continuously recorded by a computer system and uploaded to a remote server.

### RFID box use is season and sex specific

The barn provides shelter from wind and precipitation but is exposed to natural seasonal fluctuations in humidity and temperature which can range from below -10° C in winter to above 25° C in summer. As we hypothesized that these seasonal changes may be relevant for understanding the relationship between vocal communication and social structure, we first sought to understand in greater detail how wild mouse social interactions depend on time of year. To do this, we analyzed patterns of RFID box use by all mice detected in the barn during a ten-year period beginning January 1st, 2013 and ending August 31st, 2023. We then asked how these patterns changed over the course of 3-month seasonal increments in which spring corresponds to March, April, and May; summer to June, July, and August; autumn to September, October, and November; and winter of a given year to December of that year plus January and February of the next.

6,821 mice used at least one box in the barn between 2013 and 2023 (3,457 males, 3,305 females, and 59 mice with missing sex information), resulting in 39,300,829 individual box stays. During this time, an average mouse spent 55 cumulative days inside boxes during 5,654 unique stays lasting on average 3 minutes each (minimum average stay length per mouse: 0.01 seconds; maximum: 23.5 hours), all over the course of an average of 148 days from time of first detected box entrance to last detected exit (Figure 2A). A typical mouse therefore occupied the barn for a period of several months, with many days of cumulative box use consisting of box stays that tended to be short on average but were highly variable across mice. While boxes were used throughout the day, box stays peaked at dawn and dusk, corresponding to the primary hours of activity for crepuscular house mice^35^ (Figure 2B). Mice co-occupied boxes with 2.4 other individuals on average, with most of these interactions taking place in the early morning (Figure 2C). Thus, mice largely use boxes in the context of social interactions with other mice, consistent with previous work in this population focusing on their use for breeding and nesting^24,36,37^.

**Figure 2.**
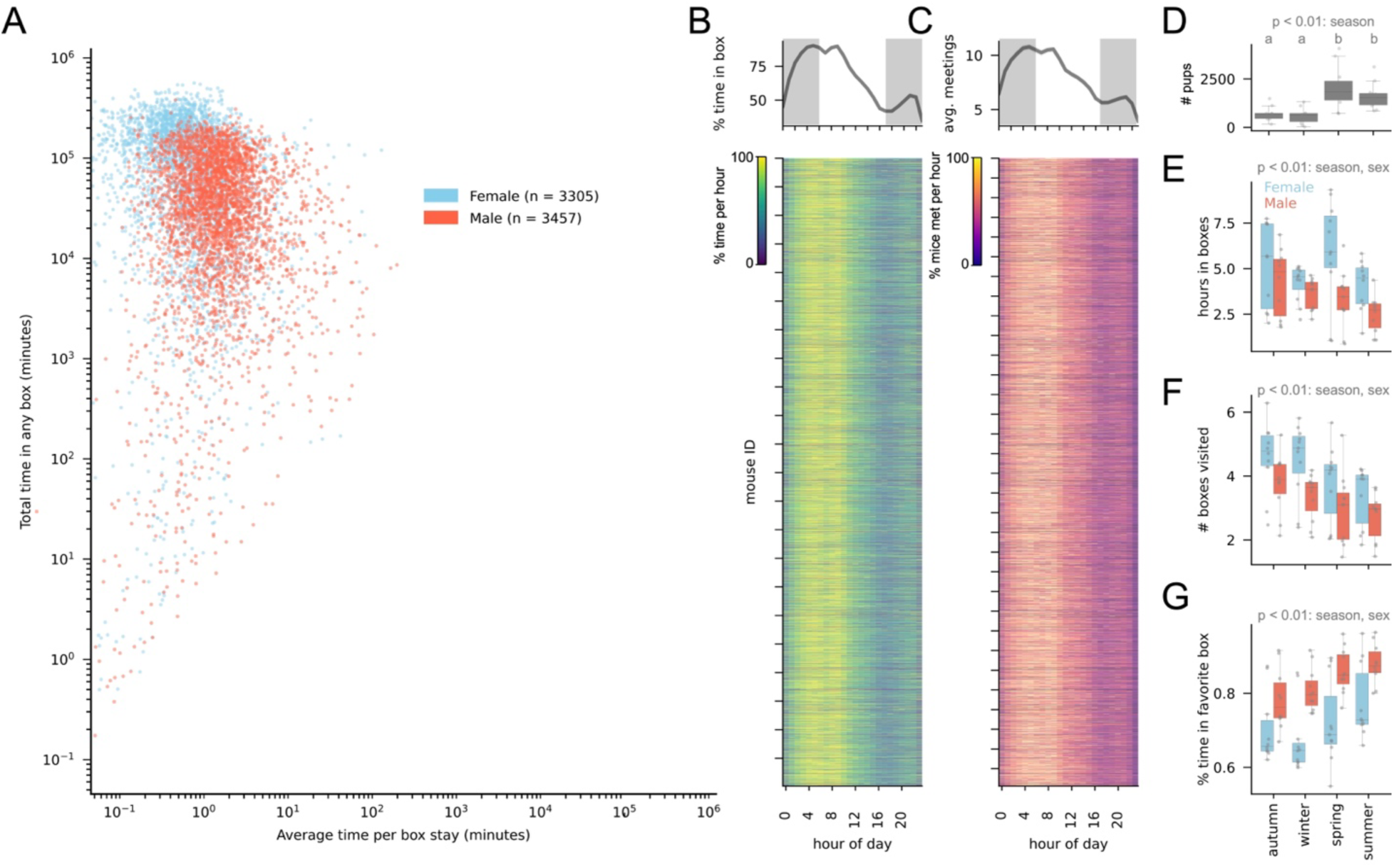
Season and sex specific patterns in RFID box use by individuals. **(A)** Total time in boxes (y axis) vs. time per stay in a box (x axis) for all mice detected in any box between 2013 and 2023. Blue: Female; Red: Male **(B)** Top: box use by hour of day for all mice detected in any box between 2013 and 2023, shading indicates nighttime hours using average yearly sunrise and sunset time for the location of the barn; Bottom: heat map of data in top plot, one row for each mouse detected in any box between 2013 and 2023. Color indicates the percent of each mouse’s time it spent inside boxes at each hour of the day, normalized to the hour with the most stay time **(C)** Top: number of meetings with other mice by time of day for all mice detected in any box between 2013 and 2023; Bottom: heatmap of data in top plot, one row for each mouse detected in any box between 2013 and 2023. Color indicates the percent of each mouse’s meetings that occurred at each hour of the day, normalized to the hour with the most meetings (**D**) Number of pups discovered per box by season for all years between 2013 and 2023, one-way ANOVA with Tukey post hoc test for the effect of season on pups discovered at each nest check during this period, p < 0.001 **(E)** Hours spent in boxes by season and sex for all mice detected in at least one box between 2013 and 2023, two-way ANOVA with Tukey post hoc test, p < 0.05. **(F)** Number of boxes visited by season and sex for all mice detected in at least one box between 2013 and 2023, two-way ANOVA with Tukey post hoc test, p < 0.01 **(G)** Percent time spent in box with most time spent (“favorite” box) by season and sex for all mice detected in at least one box between 2013 and 2023, two-way ANOVA with Tukey post hoc test, p < 0.01.

As these studies indicate that reproduction in this population is seasonal, with most pups born in spring and summer (Figure 2D; one-way ANOVA with Tukey post hoc test, p < 0.001), we next asked how these patterns of box use depended on both time of year and mouse sex. We found that females spent more time in boxes than males, an effect that was driven largely by sex differences in box use in spring and summer (Figure 2E; two-way ANOVA with Tukey post hoc test, main effect of season and sex, p < 0.05). Females also visited more unique boxes than males, and overall mice tended to visit more boxes in winter and autumn than in spring and summer (Figure 3F two-way ANOVA with Tukey post hoc test, main effect of season and sex, p < 0.01). When mice visited multiple boxes, however, they showed a preference for a single box that was stronger in males than in females, and stronger for all mice in spring and summer (Figure 2G; two-way ANOVA with Tukey post hoc test, main effect of season and sex, p < 0.01). Thus, males and females differ in their use of RFID boxes, and both change how they use boxes in a season-dependent manner.

**Figure 3.**
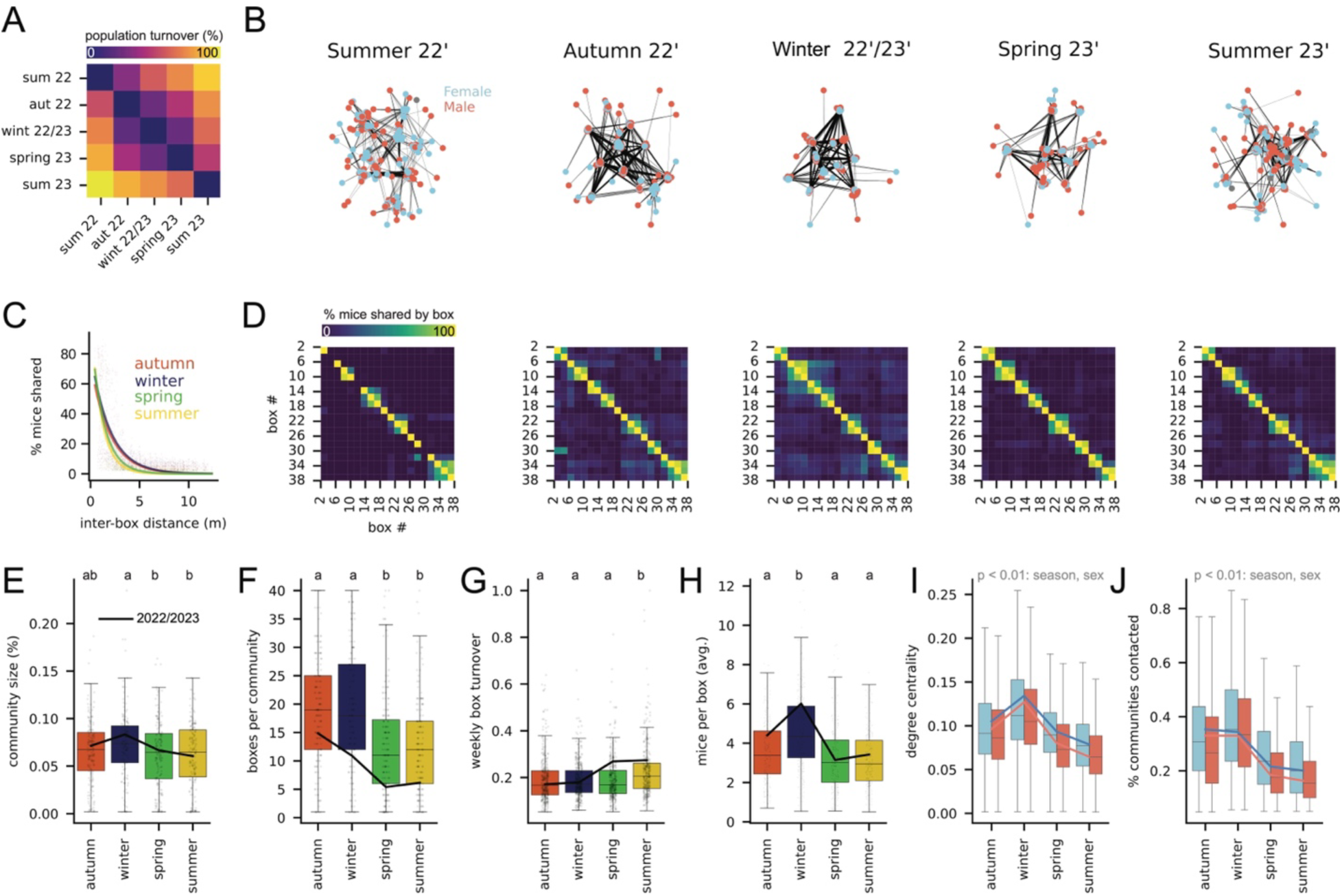
Seasonal patterns in features of social networks. **(A)** Seasonal turnover in mice by season for 2022-2023. Values represent percent of non-overlapping mice in each pair of seasonal social networks for this time period. **(B)** Social network graphs for each season between summer of 2022 and summer of 2023. **(C)** Percent mice shared between boxes by box distance for all seasons and years between 2013 and 2023. Colored lines are exponential decay models (y = a * -b^x^) fitted to data from each season, pooling across years. **(D)** Percent of mice shared between box pairs for all seasons between summer of 2022 and summer of 2023. (**D**) Community size relative to population size by season for all years between 2013 and 2023, one-way ANOVA with Tukey post hoc test, each value represents one detected community from one seasonal social network, p < 0.05. Letters indicate significantly different groups. Black line connects average values for seasons 2022-2023. **(F)** Number of unique boxes visited per community for all years between 2013 and 2023, one-way ANOVA with Tukey post hoc test, p < 0.001, each value represents total boxes visited per community per seasonal social network. Letters indicate significantly different groups. Black line connects average values for seasons 2022-2023. **(G)** Average weekly change in box occupancy for all years between 2013 and 2023, one-way ANOVA with Tukey post hoc test, each value represents the average weekly turnover of a single box per season for each year between 2013 and 2023, p < 0.001. Letters indicate significantly different groups. Black line connects average values for seasons 2022-2023. **(H)** Average mice per box for all years between 2013 and 2023, one-way ANOVA with Tukey post hoc test, p < 0.001, each value represents the average number of mice per box per season. Letters indicate significantly different groups. Black line connects average values for seasons 2022-2023. **(I)** Node centrality for all males and females in all social network graphs between 2013 and 2023, two-way ANOVA, main effect of sex, p < 0.01, and main effect of season, p < 0.01, and effect of their interaction, p < 0.01. Each value represents one individual per seasonal social network. Red line: males 2022-2023. Blue line: females 2022-2023. **(J)** Number of distinct communities contacted, where contact is defined as having spent any amount of time in a box with a mouse from a given community, by sex and season for all males and females in all social network graphs between 2013 and 2023. two-way ANOVA, main effect of sex, p < 0.01, main effect of season, p < 0.01, and effect of their interaction, p < 0.01. Each value represents one individual per seasonal social network. Red line: males 2022-2023. Blue line: females 2022-2023.

### Wild house mice occupy seasonally dynamic social groups

How are these patterns of box use by individual mice related to population-wide social structure? To address this question, we generated season-specific social networks from partially overlapping (Figure 3A) groups of mice in which nodes represent individuals and edge weight represents how much time pairs of mice spent together in any box, normalized to their total box use (Figure 3B, Supplemental Figure 2A; see methods for details). These social networks were significantly more modular than expected by chance (Supplemental Figure 2B; t-test, p< 0.01), indicating a high level of social structure. To better understand this structure, we first asked how groups of mice were distributed among boxes in the barn. We found that boxes that shared mice were typically adjacent and that the distance between these boxes was larger in winter and autumn than in spring and summer (Figure 3C,D, Supplemental Figure 2C), suggesting that groups of interacting mice occupy territories whose size expands in winter and contracts in summer.

We next partitioned graphs into communities of interacting mice (using the Louvain method; see methods section for details), then asked how features of these communities differed across seasons for the last ten years. We found that communities tended to be larger (Figure 3E; one-way ANOVA with Tukey post hoc test, p < 0.05) and occupy a larger number of boxes (Figure 3F; one-way ANOVA with Tukey post hoc test, p < 0.001), in winter than at other times of year. Consistent with these trends, boxes experienced less week-to-week turnover in their occupants (Figure 3G; one-way ANOVA with Tukey post hoc test, p < 0.001), and contained more mice (Figure 3H; one-way ANOVA with Tukey post hoc test, p < 0.001) in winter than in summer during this period. Given their differences in RFID box use, we also asked whether males and females differed in their location within seasonal social networks. Across all seasons, females interacted with more mice than males, resulting in higher network centrality (Figure 3I; two-way ANOVA, p < 0.001), and were also more likely to interact with mice from multiple communities other than their own (Figure 3J; two-way ANOVA, p < 0.001). Thus, social structure in this population exhibits seasonal dynamics in which males and females play different roles.

The barn population underwent a dramatic population decline in the autumn of 2021 following a series of cat predation events, resulting in the loss of hundreds of individuals (Figure 1F). As similar catastrophic events have been shown to restructure social networks in primate populations^38^, we considered what effect this event had on social networks in the barn. In contrast to changes in social interactions reported in primates, we found that neither the number of interactions per mouse (i.e., node density) nor the extent to which the population wide social network could be divided into communities (i.e., graph modularity) were affected by the population crash (Supplemental Figure 2D, E; t-test comparing season-specific social networks in the two years prior to the year of the crash and the two years following it, n.s.). We did find that average node clustering coefficients were higher in social networks following the crash (Supplemental Figure 2F; t-test, p < 0.01), although we cannot exclude that this is due the intentional reduction in box number that occurred in February of 2021. With this exception, however, social structure in the barn population appears to be largely resilient to large changes in population size, consistent with work on previous predation events showing little effect on social group composition^39^.

To explore what might explain this resilience, we tested if mouse social networks, like those of humans, have “small world” features, i.e. high overall clustering and short average path lengths, which have been proposed to make social networks resilient to the loss of individuals ^40^. Using a metric that compares these two features, the small-world coefficient^41^, we found that this was the case: all seasonal networks for the last 10 years had small-world coefficients significantly greater than 1 (one sample t-test, p < 0.01), indicating higher levels of clustering relative to path lengths compared to randomized control networks (see methods for details). This may explain the resilience of the barn’s social networks to both the predation events and RFID box removal that occurred in 2021. Taken together, these results describe a population of wild mice that is characterized by seasonally fluctuating small-world networks in which social group size and stability are highest in winter and autumn, lowest in spring and summer, and females play a central role in group connectivity.

### Vocal communication in wild house mice is seasonal

Acoustic communication has been proposed to play an important role in mediating group dynamics in social mammals, although a majority of studies supporting this role have focused on primates^11^. As a result, much less is known about the role that it plays in other social mammals. The seasonal social dynamics we observe in the barn population thus afforded an opportunity to ask what role acoustic communication plays in dynamic social groups of house mice, where a wealth of data already exists on neural circuits for vocalization^33^. To do this, we recorded vocalizations in RFID boxes over the course of a single seasonal cycle, beginning in August of 2022 and ending in November of 2023 (Supplemental Figure 1A). Specifically, we used AudioMoth acoustic loggers to collect two-day long audio recordings from RFID boxes approximately twice a month for 15 months, resulting in 144 recorded boxes comprising 6,594 total hours of audio (Supplemental Figure 1B-D, Supplemental Figure 3A, B). We then manually searched for and annotated mouse vocalizations in a subset of these recordings, which confirmed that AudioMoths were capable of detecting the two primary vocalization types in the house mouse vocal repertoire: low frequency squeaks and high frequency ultrasonic vocalizations, i.e. USVs (Supplemental Figure 3C, D).

We then trained and evaluated a convolutional neural network on these annotations (Supplemental Figure 4 E-K, squeak F1 score: 0.85, USV F1 score: 0.76, see methods for details), and used it to automatically identify vocalizations of each type. This revealed 1,366,171 total vocalizations, corresponding to approximately 3 vocalizations per minute from an average RFID box, with vocalizations labeled as squeaks being approximately 10 times more abundant than those labeled as USVs (1,260,452 squeaks vs 105,719 USVs). To validate these labels, we generated spectrograms for each vocalization and visualized their locations in acoustic space using UMAP (Figure 4A). Spectrograms fell into one of two acoustically clusters that with few exceptions corresponded to predicted class labels (Figure 4B).

**Figure 4.**
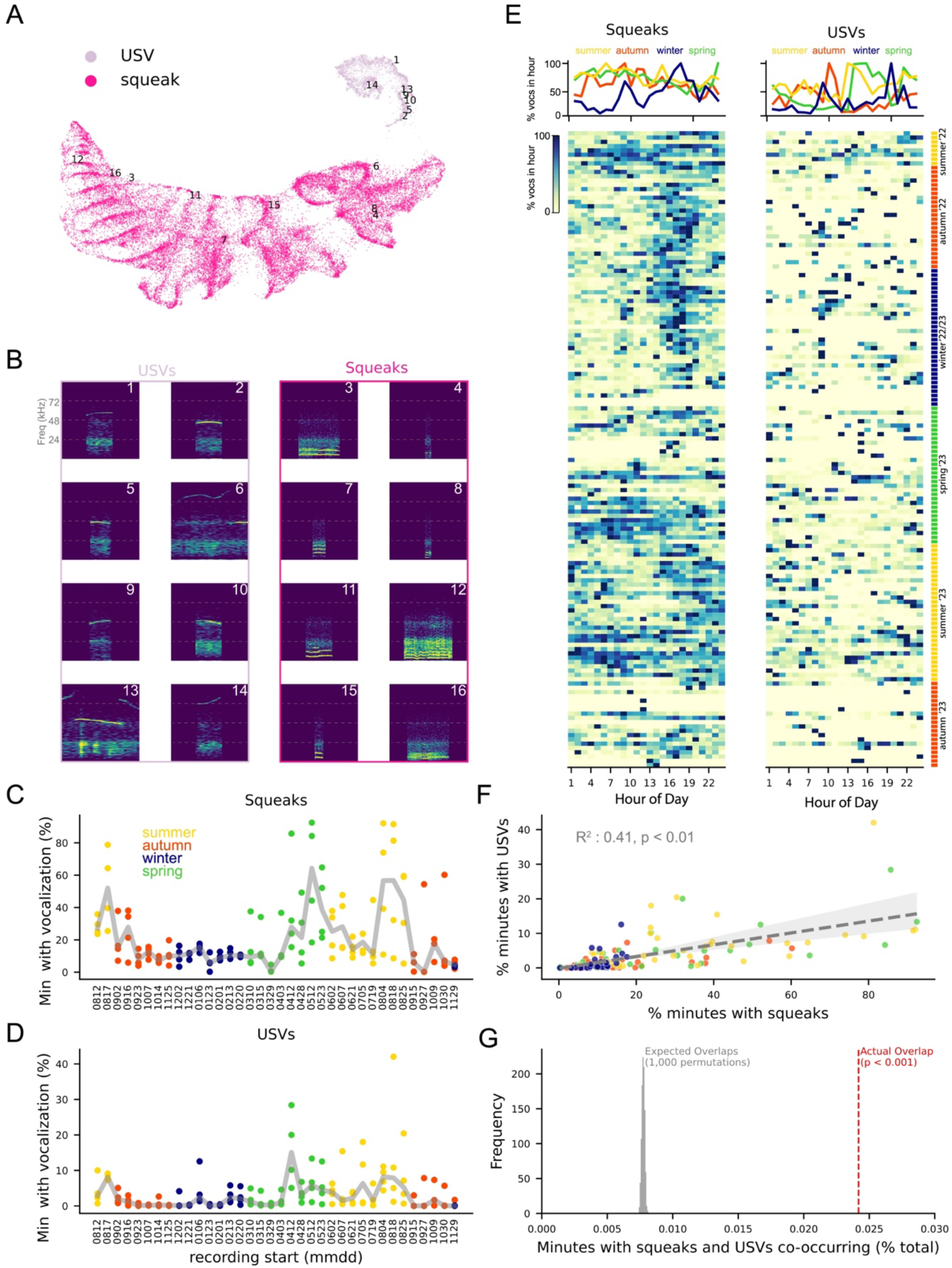
Vocal communication is seasonal in wild house mice. **(A)** UMAP embedding of 225,334 spectrograms sampled randomly from detected squeaks and USVs. Each dot corresponds to a single spectrogram representing one vocalization. Numbers on representative dots correspond to spectrograms in panel B. **(B)** Example spectrograms from panel A. **(C)** Percent minutes with squeaks vs recording start date, one-way ANOVA with Tukey post hoc test comparing percent minutes with squeaks by season, p < 0.001 **(D)** Percent minutes with USVs vs recording start date, one-way ANOVA with Tukey post hoc test comparing percent minutes with USV by season, p < 0.001 **(E)** Number of squeaks (left) and USVs (right) by time of day. Top: Average percent of vocalizations per hour by season across all recorded boxes. Bottom: heatmaps of data in top plot, each row corresponds to a single RFID box recording and color corresponds to the percent of all vocalizations in a given recording falling in that hour, normalized to the hour with the most vocalization. **(F)** Squeaks per recording vs USVs per recording with linear fit, R^2^= 0.41, p < 0.01, shading above and below line corresponds to the 95% confidence interval **(G)** Actual percent of minutes containing both squeaks and USVs (red line) and percent of minutes containing both squeaks and USVs from 1,000 permutations of squeak timestamps relative to USV timestamps (gray), p < 0.01.

To begin characterizing the relationship between these vocalizations and seasonal changes in social groups, we first asked if the amount of time mice spent vocalizing differed between recording dates. We found that it did for both squeaks and USVs and that these differences were significantly affected by the season in which the recording was made (Figure 4C, D; one-way ANOVA with Tukey post hoc test, effect of season on squeaks: p < 0.001; effect of season on USV: p < 0.001), with the majority of minutes containing vocalization (87.8%) occurring in the summer and spring compared to winter and autumn. Thus, vocal communication is seasonal in this population.

As we observe daily rhythms in box use by individual mice (Figure 2B), we also asked if the number of vocalizations produced in recorded boxes depended on time of the day. This was true for squeaks, but only in winter when they were most abundant shortly before and after sunset (Figure 4E, left; one-way ANOVA with Tukey post hoc test comparing vocalization count by hour of the day, p < 0.001). Vocalizations in autumn exhibited a qualitatively similar diurnal pattern to those observed in winter, but this effect was not statistically significant, nor were qualitative patterns of increased vocalization in early morning and evening during spring and summer (Figure 4E, left; one-way ANOVA with Tukey post hoc test, n.s.). We did not observe a significant effect of time of day on USV production during any season (Figure 4E, right).

Squeaks and USVs have been proposed to signal distinct affective states in house mice ^42^, suggesting they may be used in different social contexts. To test this prediction in wild mice, we asked how often squeaks and USVs were detected in the same recorded minute in our dataset and compared this to the null hypothesis that production of each call type was independent of the other. We found that amount of time mice spent producing USVs in a given box was positively correlated with that for squeaks (Figure 4F, R-squared = 0.414, p < 0.01), and that across seasons the number of minutes containing both vocalization types was significantly higher than expected by chance (Figure 4G, p < 0.001). Thus, although squeaks and USVs may signal different kinds of social information, wild mice appear to produce them in close temporal association in at least some social contexts.

### Vocal communication is associated with the presence of pups

These findings suggest a broad correlation between vocal communication and seasonal patterns in mouse social structure: mice vocalized least at the times of year when they occupied the largest and most stable social groups (i.e. autumn and winter) and most at the times of year when those groups were smallest and most dynamic (i.e. spring and summer). Nonetheless we find the vocal communication is highly variable from box to box even when those boxes were recorded simultaneously. To identify sources of this variation, we next asked what features of social groups predicted the amount of vocalization they produced.

To do this, we considered four hypotheses about the relationship between group composition and vocalization, and compared them using the Akaike Information Criterion (AIC^43^): first, that vocalization is seasonal but not otherwise related to box occupants (vocalization ∼ season); second, that vocalization is seasonal and also depends on the number of mice using the box over the course of the recording (vocalization ∼ season*# mice); third, that it depends on the proportion of males to females using the box rather than total number of mice (vocalization ∼ season*proportion male); and fourth, that it depends on the number of pups discovered in the box at the end of the audio recording (vocalization ∼ season*pups).

The model in which vocalization depends on season and number of pups had the lowest AIC score for both squeaks and USVs, and, for squeaks, also explained the largest percent of the variance in vocalization counts (Figure 5A, Table 1). Consistent with this assessment, the average number of squeaks and USVs per deployment closely tracked the number of pups discovered in each recorded RFID-box (Figure 5B) during each of the two waves of pups that occurred in 2023, one in early spring and one in late summer. Thus, on the timescale of days, the number of pups, more so than the number of adults or their sex ratio, is predictive of how much vocalization occurs in RFID boxes.

**Figure 5.**
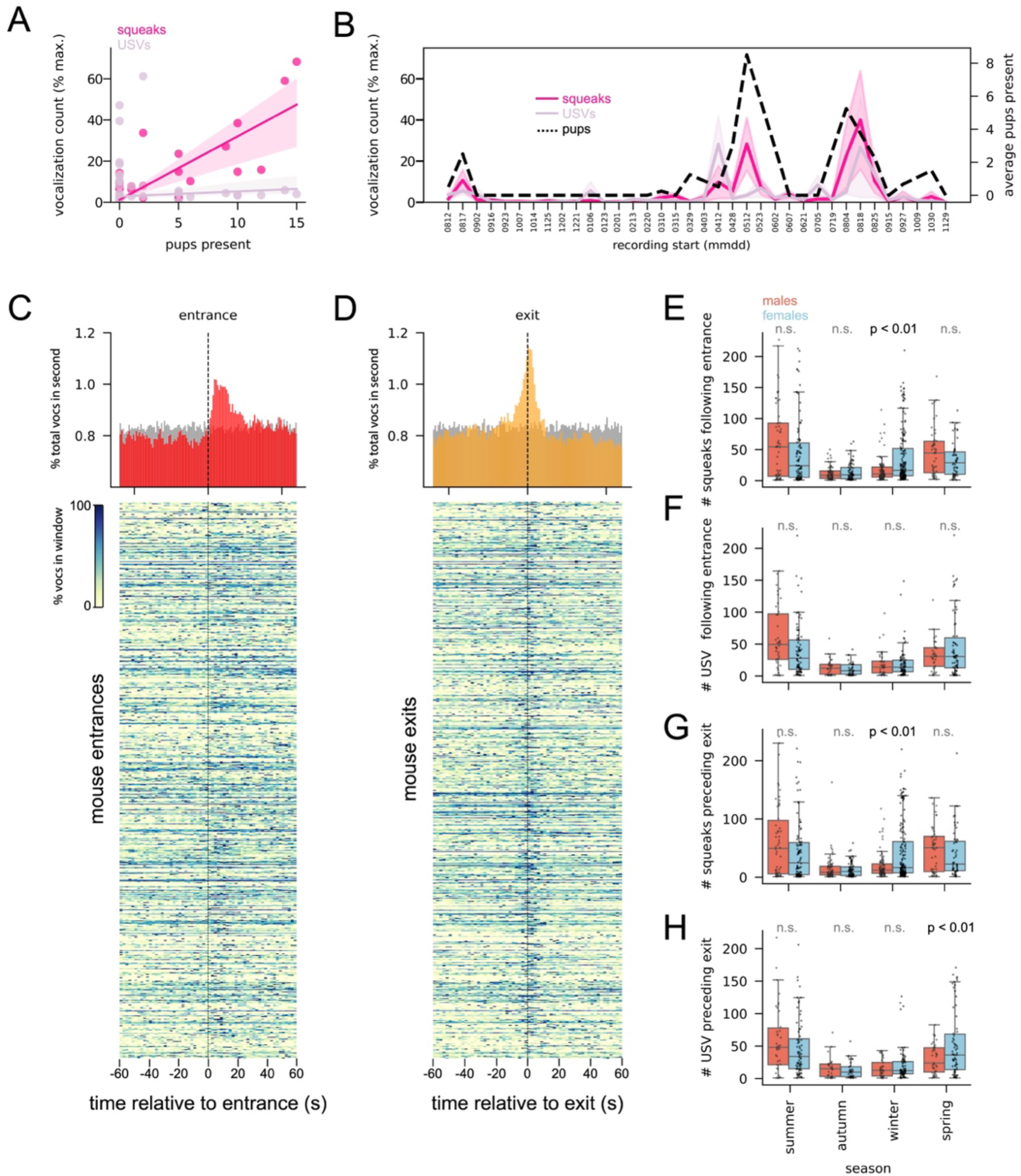
Vocalization is associated with pups and changes in RFID box occupancy. **(A)** Linear fit of pup number present at the end of each recording by squeaks (magenta) and USVs (light pink) detected in that recording. Shading above and below each line corresponds to the 95% confidence interval. **(B)** Average squeaks and USVs recorded and average pups detected in all recorded boxes by recording date. Lines represent average per recording date. Shading represents 95% confidence interval. Vocalization counts are the same used to generate plots in figure 4 **(C)** Top: percent of all vocalizations aligned to box entrance in one second bins, for all box entrances containing at least one vocalization within 1 minute preceding or following the entrance. Bottom: data in top histogram by entrance, with one entrance per row, in 2 second time window bins. Color corresponds to percent of vocalizations per bin relative to bin with the most vocalizations. **(D)** Top: all vocalizations aligned to box exits, for all box exits containing at least one vocalization within 1 minute preceding or following the exit. Bottom: data in top histogram by entrance, with one exit per row, in 2 second time window bins. Color corresponds to percent of vocalizations per bin relative to bin with the most vocalizations. **(E)** Average number of squeaks within one minute following individual mouse entrances by sex and season; p-values correspond to t-tests comparing sex within each season following Bonferroni correction (**D**) Average number of USVs within one minute following individual mouse entrances by sex and season; p-values correspond to t-tests comparing sex within each season following Bonferroni correction **(G)** Average number of squeaks within one minute preceding individual mouse exits by sex and season; p-values correspond to t-tests comparing sex within each season following Bonferroni correction **(H)** Average number of USVs within one minute preceding individual mouse exits by sex and season; p-values correspond to t-tests comparing sex within each season following Bonferroni correction.

**Table 1.**
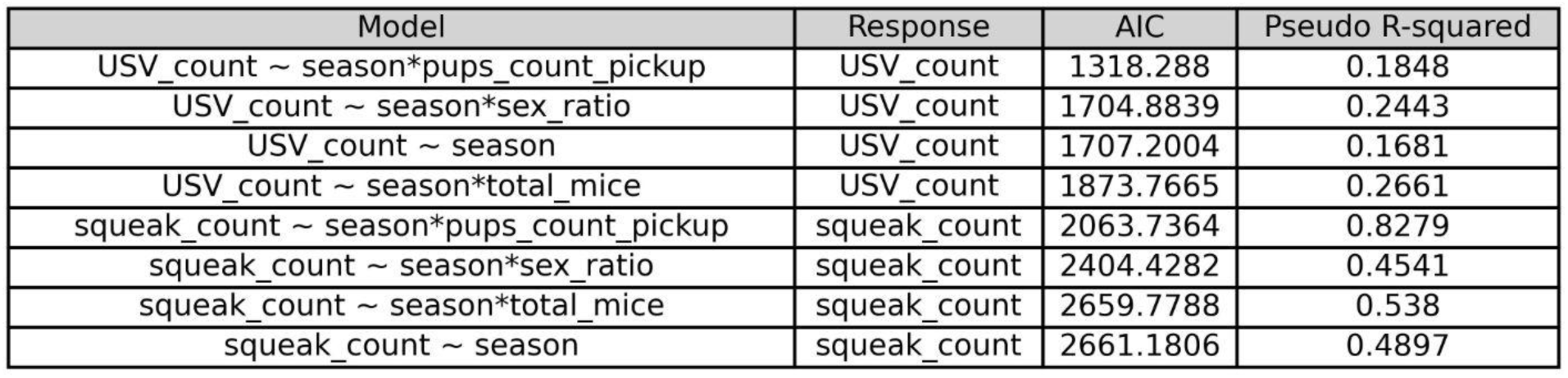
AIC and Pseudo R-squared scores for models compared in Figure 5. Variable names: USV_count: the total number of USVs detected during the recording; season: the time of year of the recording with spring corresponding to recordings taken during March, April, and May, summer corresponding to recordings taken during June, July, and August; autumn corresponding to recordings taken during September, October, and November, and winter corresponding to recordings taken during December, January, and February; total_mice: the total number of unique mice using the box during the recording; sex_ratio: the ratio of males to females using the box during the recording; squeak_count: the total number of squeaks detected during the recording; pups_count_pickup: the number of pups observed in the box at the end of the recording

### Vocal communication is associated with box entrances and exits and is correlated with the amount of time mice spend together

These analyses focused on total vocalizations emitted during 2-day long recordings, and thus leave open the question of whether vocalization is associated with individual social interactions that occur on shorter time scales. While we did not collect video data to directly link such interactions with vocalization, RFID-generated timestamps allowed us to ask if vocalization was associated with events that change social group composition, i.e. individual entrances and exits from boxes. We found that 42% of all entrance and exit events contained at least one vocalization within this window (12.2% contained a USV; 39.9% contained a squeak), and that vocalizations were closely associated in time with these events (Figure 5C, D, red and orange), trends we did not observe in simulated datasets in which vocalization times were randomized relative to entrance and exit times (Figures 5C and D, gray). The amount of vocalization surrounding box events was also affected by both season and sex of the mouse exiting or entering. Overall, more vocalizations surrounded entrances and exits in spring and summer, consistent with these seasons containing more vocalization (one ANOVA with Tukey post-hoc test, p < 0.001 for main effect of season), and more vocalization surrounded entrances and exits of females compared to males, with the largest difference between sexes occurring in winter (Figure 5E-H, Bonferroni corrected t-tests comparing sex within season).

Taken together, the above analyses indicate the vocal communication in wild house mice is correlated with features of social groups at both long (seasonal) and short time (seconds) scales. If vocal signals mediate social dynamics, however, they should also be correlated with the strength of social interactions between individual pairs of mice. To test this prediction, we calculated correlation coefficient between two time series for each unique pair of mice that spent time together in a recorded box: first, the number of vocalizations they experienced at a given meeting and second, the amount of time they had spent together by the start of their next meeting (Figure 6A). We found that vocalization was overall positively correlated with the amount of time pairs spent together for both squeaks and USVs (Figure 6B, magenta/pink: actual correlations; gray: correlations from shuffled data; t-test comparing actual to shuffled data, p < 0.001 for both vocalization types), and that these correlations were season and sex dependent. Specifically, the amount of time females spent together was more positively correlated with vocalization than males for both squeaks and USVs, and this was due to differences between sexes in the spring, when the amount of time that male-male pairs spent together was more likely to be negatively correlated with vocalization than that of female-female pairs (Figure 6C; see figure legend for details).

**Figure 6.**
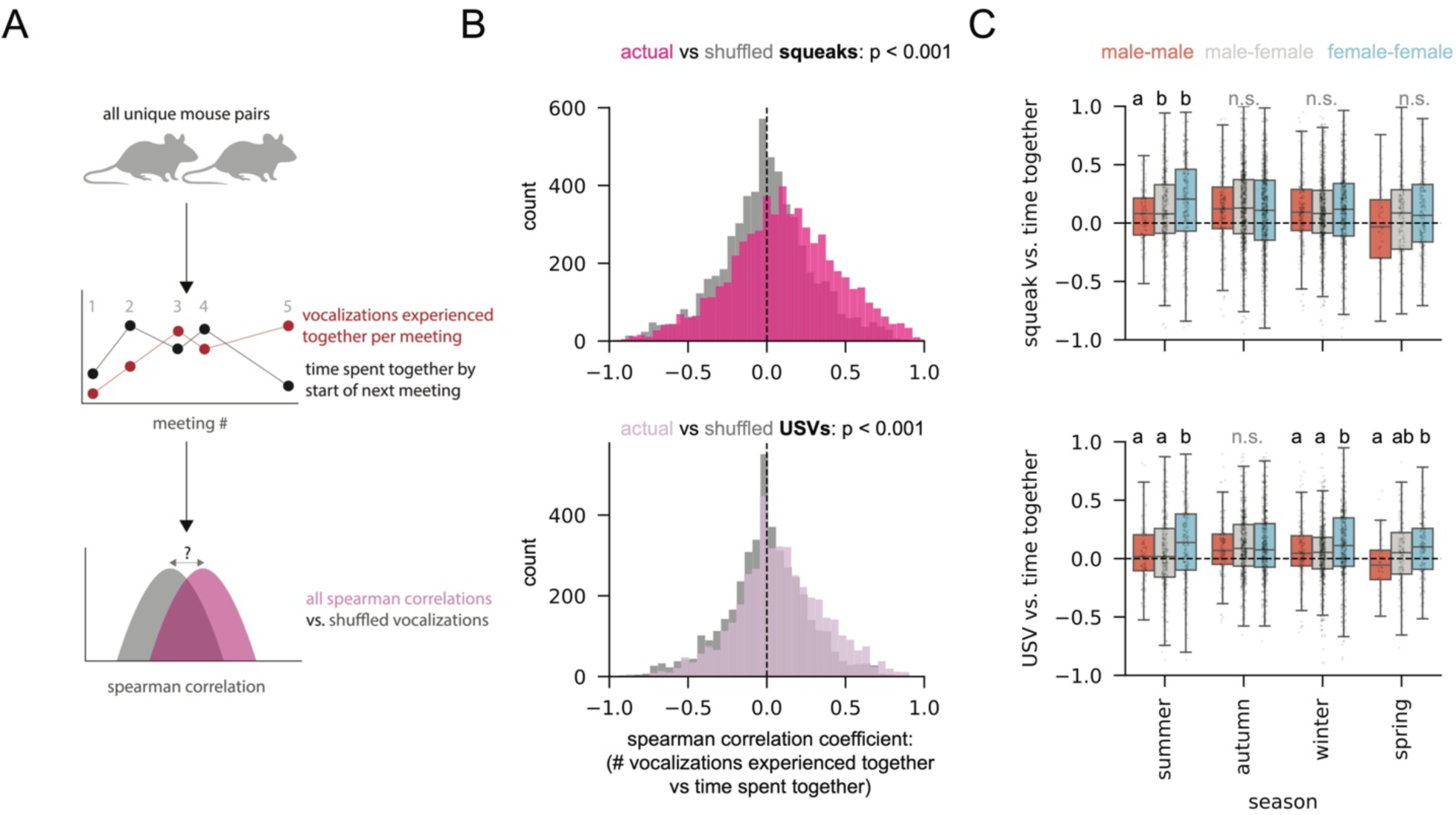
Vocalization is correlated with how much time mice spend together. **(A)** Schematic illustrating analyses in panels B and C: for each unique pair of mice, Spearman correlation coefficients were calculated between vocalization experienced together in each meeting and time spent together by the start of the next meeting (example cartoon shows one mouse pair meeting 5 times). Distributions of these correlations coefficients were then compared to coefficients generated from the same analysis with vocalization counts shuffled relative to time spent together **(B)** Distributions of Spearman correlation coefficients between squeaks and time spent together for squeaks (magenta, top) and USVs (pink, bottom), compared to shuffled controls (gray). Actual distributions differed significantly from controls for both vocalization types (t-tests, p < 0.001). **(C)** Top: The same correlation coefficients in panel B, top, split by sex and season. Main effect of season, p < 0.01, Interaction between sex and season p < 0.01, no effect of sex. Two-way ANOVA. Letters indicate significantly different groups in within-season ANOVA. Bottom: The same correlation coefficients in panel B, bottom, split by sex and season. Main effect of season, p < 0.01, Interaction between sex and season p < 0.01, main effect of sex, p < 0.01. Two-way ANOVA. Letters indicate significantly different groups in within-season ANOVA.

## Discussion

Using automated behavior tracking, we characterize social interactions in a population of wild house mice over the course of ten years. Consistent with previous work examining shorter time scales^34^, we find that mice in this population cluster into social groups, that these groups are seasonal, and that female mice play an important role in their organization. In addition, we find that the use of vocalization in this population is seasonal, correlated with changes in group composition following box entrances and exits, and most likely to take place in the spring and summer when social groups were smallest, most dynamic, and contained pups. Taken together, these results provide insight into the social structure of wild mouse populations and the potential roles that vocal communication may play in contributing to that structure.

Our approach has the advantage of investigating behavior in free-living mice exposed to natural variation in environmental variables such as weather and temperature. As a result, behavioral variance in our dataset is high. A complementary strategy is to focus on small populations over short timescales during which environmental variables are relatively stable. For example, recent studies have used RFID tracking in tens of rewilded laboratory mice exploring outdoor enclosures over periods of several days during summer months. In these short-term experiments, male mice quickly established and defended territories at the exclusion of other males, and strain-specific female behaviors drove population-wide social structure^25,44^.

Our findings in wild mice are generally consistent with these results. For example, males in our study population exhibited stronger box preferences than females, suggesting the establishment of male-specific territories (although multiple males do co-inhabit RFID boxes), and overall females were both more central in social networks than males and interacted with a larger number of distinct social groups, suggesting a larger role in shaping the structure of population wide social networks. However, monitoring large populations in natural environments over long time scales further allowed us to observe an effect of season on these behaviors. While males exhibited stronger box preferences than females across all seasons, all mice had stronger preferences in summer and spring compared to autumn and winter. In addition, females spent more time using RFID boxes overall than males, a difference that was most dramatic in spring. We hypothesize that seasonal changes in both temperature and sex-specific reproductive behaviors shape these behavioral patterns, and future work will use long term environmental datasets collected at the barn to disambiguate the relative contributions of these environmental and biological variables to seasonal social dynamics.

Consistent with laboratory studies of house mouse vocalization, we find that the vocalizations produced by wild house mice fall into one of two categories: ultrasonic vocalizations (USVs) and low-frequency squeaks. While most laboratory studies have focused on USVs, we find that squeaks outnumber USVs by a factor of ten in our dataset. This difference is probably an overestimate due to the lower sensitivity of AudioMoths at high frequencies^14^, but squeaks nonetheless appear to comprise a large proportion of the vocalizations produced by mice in wild populations. In addition, we find that they occur in close temporal association with USVs more often than expected by chance. Although we do not know the identity of vocalizing mice with boxes, we hypothesize that these associations arise from social contexts in which squeaks and USVs both convey meaningful social information. For example, squeaks and USVs have been reported to occur in close temporal association in late stages of copulation^45^, and classical work has described low-frequency “wriggling” calls produced by non-nursing pups that may occur close to isolation induced pup USVs^28^.

Tracking vocal behaviors over long time scales further revealed that they are seasonal, with the majority of vocalization occurring in the spring and summer when social groups are smallest. Thus, social vocalization in wild mice is not positively correlated with group size. Rather, we find that the presence of pups is a strong predictor of vocal communication. While we are unable to identify individual vocalizers in our recordings, a previous study which collected long term audio from families of Mongolian Gerbils found that removal of pups resulted in a decrease in vocalization^46^, suggesting that pups, either by producing vocalizations or triggering them in others, are an important source of vocal communication in rodents.

Two key limitations of our study are that we can’t align vocal events to specific social behaviors using video data, and we can’t experimentally test the effect of specific vocalizations on social dynamics. Nonetheless, we do observe a close temporal association between vocalization and events (box entrances and exits) that change group membership. In addition, we find that on average the amount of time pairs of mice spend together is positively correlated with how many vocalizations they experience together, and that these correlations depend on season and sex. We hypothesize that these sex differences result from sexually dimorphic roles for acoustic communication in house mice. For example, males may be more likely to use acoustic communication in the context of competition (e.g., for access to females in an RFID box) while females are more likely to vocalize within groups of other females with whom they are rearing pups^47^. Future studies will use acoustic playback experiments in wild populations to directly test the relationships between vocal signals and social group dynamics.

Short-term studies in the lab and in outdoor enclosures are essential to understand the genetic and neurobiological underpinnings of social behaviors. However, the amount of experimental control in these studies also limits their scope to describe and understand social behaviors in the natural contexts in which they evolved to function. Technologies for long-term, passive monitoring of social behaviors in the wild now make it possible to capture aspects of sociality that cannot be observed in the laboratory (e.g., the effect of catastrophic environmental disasters^38^). Using these tools in wild mice, this work lays a foundation to understand how behavioral variation interacts with the environment to influence sociality in the wild.

## Methods

### Data Collection

#### Study Population

We studied a population of wild house mice living in a 72 m^2^ barn located in mixed woodland and agricultural fields approximately 20 km from Zürich, Switzerland^48^. Mice were free to enter and exit the barn and were supplied with *ad libitum* food (1:1 mix of ‘‘Hafer flockiert’’, UFA AG, 3360 Herzogenbuchsee, Switzerland and ‘‘Meerschwienchen und Hamster Futter’’, Landi Schweiz AG 3293 Dotzigen, Switzerland) and water. The barn contained 20 boxes (“RFID boxes”, 15 cm tall, 15 cm diameter) with a single entrance tunnel that mice could use as they chose (prior to 2021, there were 40 boxes). Straw and soft bedding in these boxes was replaced every two months. *Mus musculus domesticus* was the only rodent species observed in the barn during the study period. Audio recordings of mice were approved under permit ZH076/2022 granted by the Veterinary Office of Canton, Zürich, Switzerland. Whole population censuses and RFID tracking between 2013 and 2023 were approved under permits ZH051/2010, ZH056/2013, ZH091/2016, ZH098/2019 granted by the Veterinary Office of Canton, Zürich, Switzerland.

#### Whole population censuses

We carried out whole population censuses of the barn population approximately once every two months. During these censuses, every mouse in the barn was caught, individually weighed, and checked for the presence of an RFID tag. Mice > 17.5g that did not have a tag were injected subcutaneously with one. After tagging, all mice were returned to the approximate location of the box from which they were caught.

#### RFID tracking

Box use was passively monitored using a custom RFID system^48^. Briefly, the circular entrance tunnel of each RFID box was fitted with two RFID readers separated by a distance of approximately 10 cm and connected by cables to a central computer inside the barn. RFID detections were logged continuously by this computer, then transmitted every 48 hours to a remote server, where they were automatically checked for errors, and saved in an SQL database.

#### Audio Recording

Recordings were made using AudioMoths (hardware version 1.2.0, firmware version 1.8.0) deployed 36 times between August 2022 and November 2023, with 2 - 3 deployments per month with the exception of November of 2022 and 2023 during which there was only one deployment. During each deployment, 4 AudioMoths were left on top of haphazardly chosen RFID boxes (one AudioMoth per box) and allowed to record with the following parameters:

Sampling Rate: 192000 Hz
Duty Cycle: 55 s recording / 5 s off
Filtering: 5 kHz high pass filter
Gain: “Medium”

Recordings lasted for 48-51 hours, after which AudioMoths were either collected and switched off or recordings ended because the onboard 64 GB microSD card became full. To protect AudioMoths from high humidity, they were placed in small plastic cases (IKEA Pruta food containers set 9×9×4cm, 150ml) in which two holes were made: one to control the DEFAULT - USB/OFF - CUSTOM switch (diameter 3.16 mm) and one for the AudioMoth MEMS microphone (diameter 1.58 mm). AudioMoths were secured inside these food containers with pieces of foam and a 2 cm x 2cm sachet of desiccant (cat litter). The hole allowing access to the switch was covered by a small piece of duct tape during recording. To further protect AudioMoth containers from chewing mice during recordings, food container cases were placed inside secondary protective boxes fitted on top of a 1 cm thick, 500 cm diameter plastic base. A 9.5 mm diameter hole was drilled in the center of the base to provide the microphone access to the RFID box.

At the start of each deployment, AudioMoths were configured either on a personal laptop (until January 2023) or on the computer receiving data from the barn’s RFID system (after first deployment of January 2023). They were then switched on (CUSTOM) and placed inside their cases. Cases were then placed inside protective containers. For each RFID box, the ceramic tile lid was removed and replaced with the protective container with the recording AudioMoth, during which time the mouse occupants were quickly noted. The barn was undisturbed by humans until the end of the deployment when the AudioMoth was removed, contents of the box noted, and ceramic lid replaced.

#### Aligning AudioMoth and RFID system clocks

We used an acoustic chime to account for differences between the onboard clock of each AudioMoth and the RFID system. Within the first 5 minutes of recording for each AudioMoth, we passed a specially designated test transponder through the outer RFID antenna of the box being recorded. The ID of this test transponder triggered an audible chime generated by the computer system in the barn, which was amplified by a Bluetooth speaker sitting directly on top of the AudioMoth. In this way the test transponder simultaneously generated a timestamp logged by the barn RFID system and a sound event (the chime) recorded and given a timestamp by the AudioMoth. This process was repeated at the end of each deployment (“recovery”), within 5 minutes of the end of each AudioMoth’s recording time. After data collection, we manually annotated the chimes from each AudioMoth at the beginning and end of each deployment, identified their corresponding timestamps in the RFID database, then adjusted the timestamps for each AudioMoth according to the deviation between these two, assuming a linear drift of the AudioMoth clock relative to the RFID clock over the course of the 48-hour deployment. For analyses relating RFID and audio data, only recordings in which clocks could be aligned in this way were used.

### Data Analysis

#### Acoustic Segmentation

We used the software package deep audio segmenter (DAS) to identify vocalizations in raw wav files recorded by AudioMoths^16^.

#### Annotation

We performed annotations in two phases. In the first phase, we calculated power spectral density in the 65-75 kHz range for a subset of recordings, then visually inspected 55 s long wav files for the presence of vocalizations. This indicated the presence of both low frequency, audible vocalizations (“squeaks”) and ultrasonic vocalizations (USVs). In the second phase, we manually annotated 2,137 squeaks (197 seconds) and 2,172 USVs (112 seconds) recorded during the summer, autumn, and winter of 2022. using the following criteria: (1) the vocalization had to be visible to the annotator in the spectrogram image of the audio (2) to be considered squeaks, vocalizations needed have a dominant frequency below 20 kHz (3) to be considered USVs, vocalizations had to have a minimum frequency above 40 kHz.

#### Neural Network Training

We used our annotations to train a convolutional neural network using DAS with the following parameters:

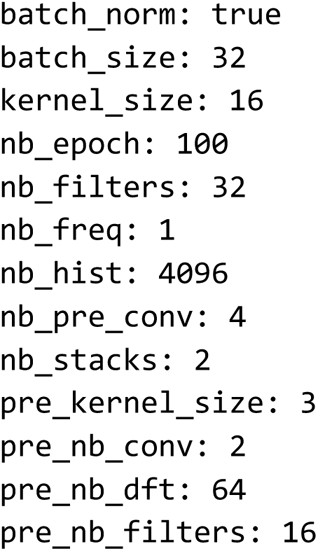

#### Neural Network Evaluation

We evaluated the performance of trained DAS networks using F1 scores calculated on withheld spectrogram chunks of the same length used during training. This calculation was implemented in DAS:

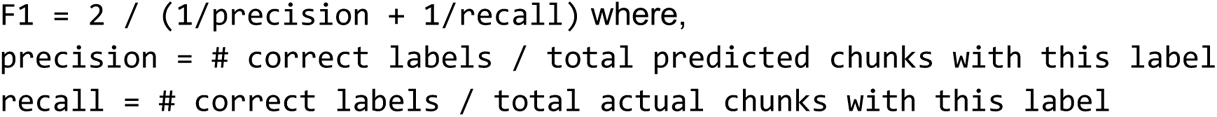

#### Inference

We performed inference using the das.predict.predict function, assigning probabilities for each class label to chunks of audio 4096 samples in length (21.3 ms). The label with the highest probability was assigned to each chunk unless no label achieved a probability higher than 0.8 (segment_thresh = 0.8). Labels separated by fewer than 5 ms were then merged into single vocalizations (segment_fillgap = 0.005). This value was chosen on the basis of inter-syllable intervals observed in annotated data. Default values were used for all other arguments of the predict function.

#### Social network construction

We constructed social networks from pairwise interactions between mice, where “interaction” is defined as spending time together inside a single box. The amount of time spent together for each pair was normalized by the total time that each mouse in the pair spent in any box without the other, i.e.

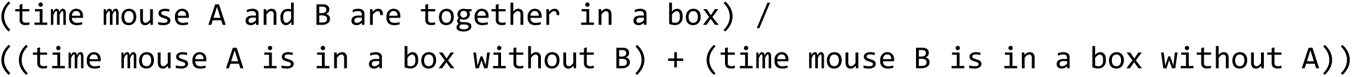

These values were calculated in 3-month seasonal intervals between 2013 and 2023, then used to generate social network graphs in igraph (https://python.igraph.org/en/stable/). Communities in these graphs were identified using the Louvain method as implemented by the best_partition method of the python-louvain package (https://python-louvain.readthedocs.io/en/latest/api.html).

#### Calculation of social network metrics

To calculate social network modularities in Supplemental Figure 2B and E, we used the modularity function of the Python package python-louvain with default values and compared modularities of actual season-specific networks to those with identical degree distributions but random connections between nodes. To calculate density and clustering coefficients in Supplemental Figures 2D and F, we used the density and average_clustering functions of networkx with default values, respectively. To determine if social networks have small world properties, we calculated a small world coefficient defined as

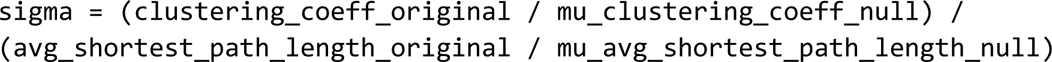

Where

`clustering_coeff_original` is the clustering coefficient of the actual network, calculated using the average_clustering function of networkx and default values,

`avg_shortest_path_length_original` is the average shortest path length of the actual network, calculated using the average_shortest_path_length function of networkx and default values

`mu_clustering_coeff_null` is the average clustering coefficient of 100 networks with the same degree distribution as the actual network, but randomized connections

`mu_avg_shortest_path_length_null` is the average shortest path length of 100 networks with the same degree distribution as the actual network, but randomized connections

To calculate degree centrality for individuals by sex in Figure 3I, we used the degree_centrality function of the community module in the Python package networkx with default values (https://pypi.org/project/networkx/).

#### Statistical Modeling

Vocalization count models, AIC values, and pseudo R-squared values in Figure 5 and Table 1 were generated using the glm module of Python package statsmodels. Data were modeled using a negative binomial distribution as our response variables were overdispersed. For each model, an alpha parameter was chosen that minimized the model’s negative log likelihood.

---

**Supplemental Figure 1.**
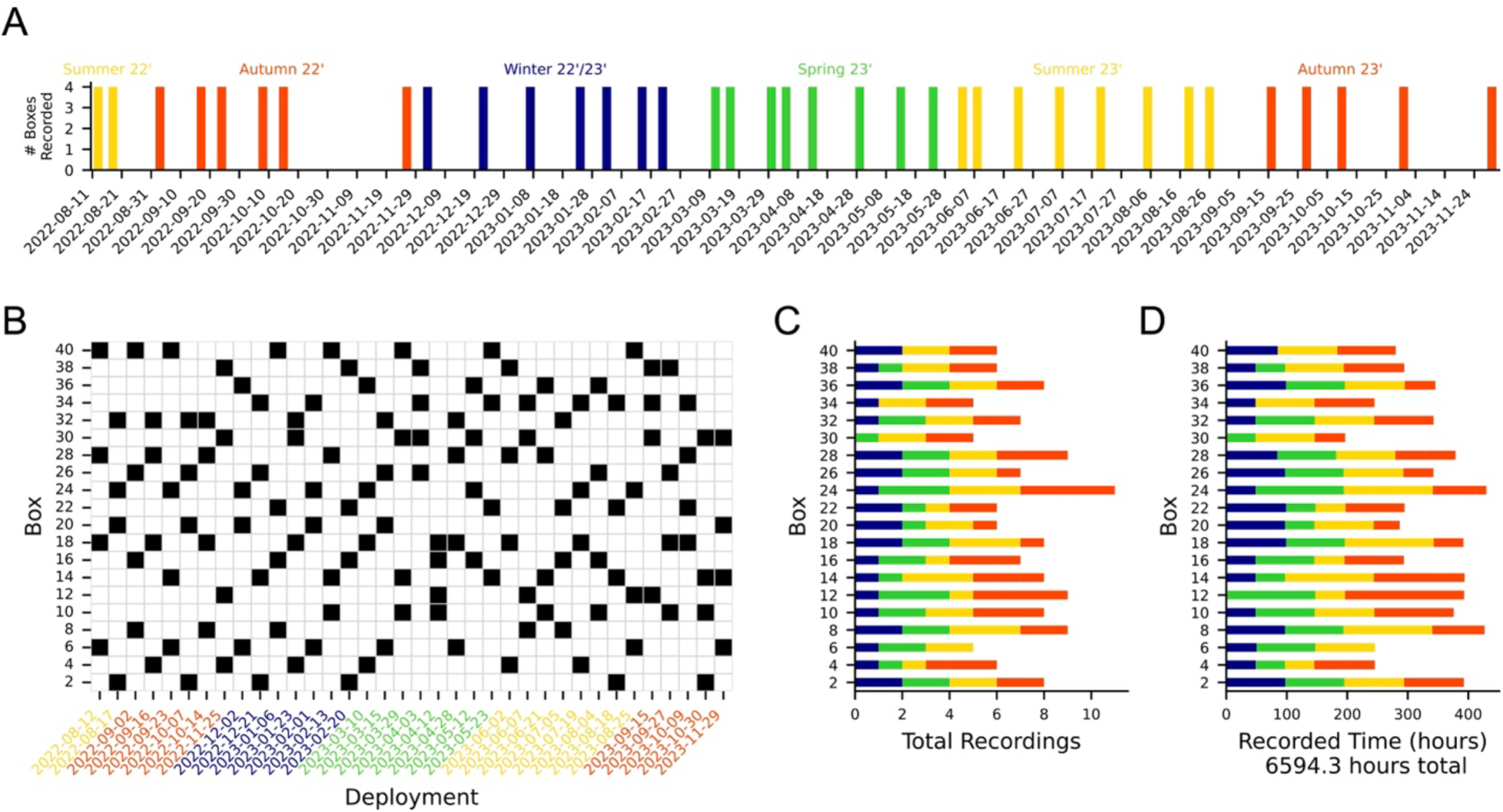
Audio Recording Regime. **(A)** Audio recording dates and number of RFID boxes recorded and analyzed per date (4 boxes for all) **(B)** Box IDs recorded at each date. See Figure 1D for box locations. **(C)** Number of recordings by box and season **(D)** Total amount of recorded audio by box and season.

**Supplemental Figure 2.**
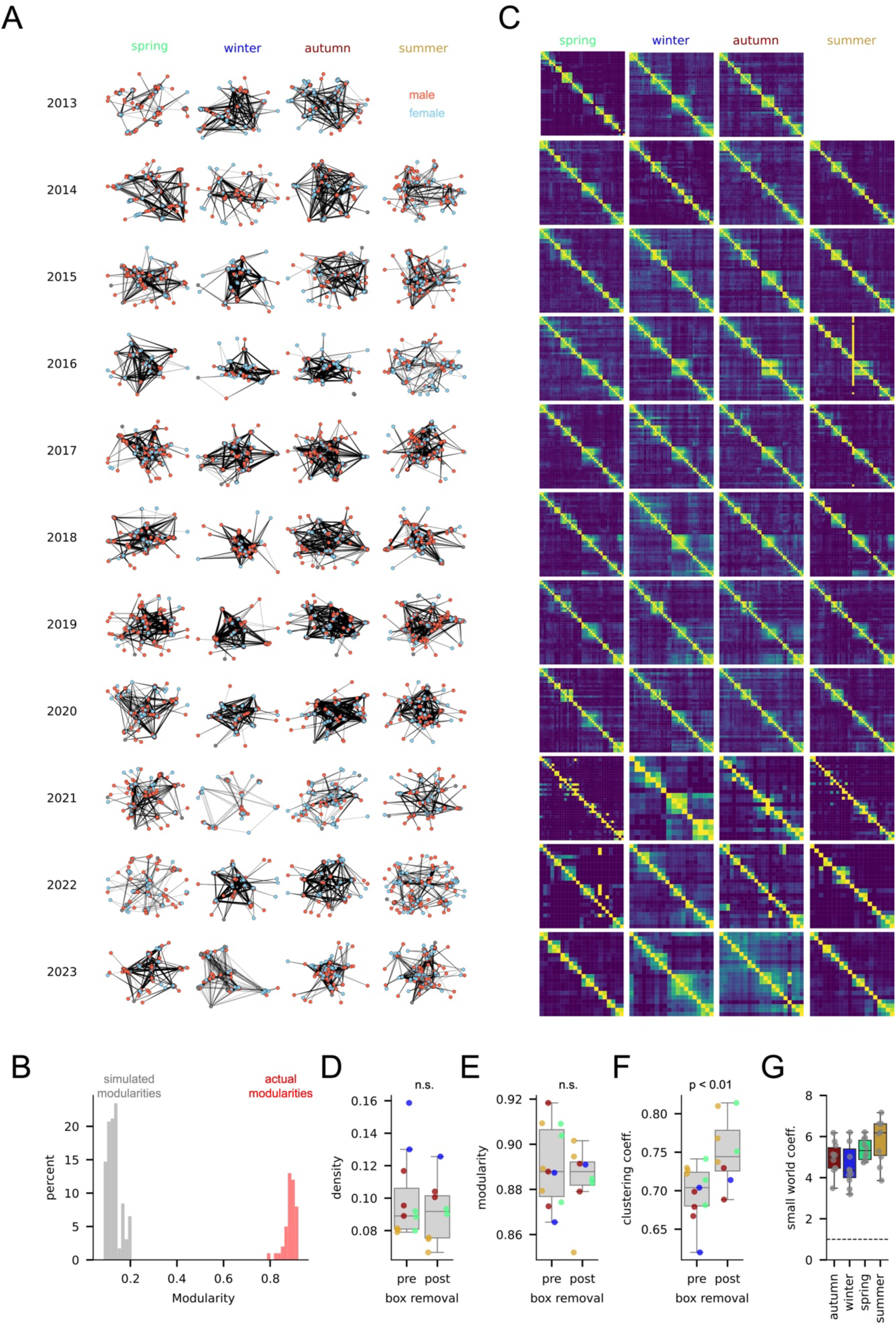
Social networks are modular and have small world features. **(A)** Social network graphs for each season and year between 2013 and 2023. Red: Male, Blue: Female. **(B)** Actual modularities of all social networks in panel A (red) compared to modularities from random graphs with the same node degree distributions (gray). **(C)** Percent of mice shared between box pairs for each year and season between 2013 and 2023. **(D)** Seasonal graph density in the 2 years pre and post removal of 20 RFID boxes, excluding seasons from the 2021 population crash, t-test, n.s. Colors correspond to seasons in panel A. **(E)** Seasonal graph modularity in the 2 years pre and post removal of 20 RFID boxes, excluding seasons from the 2021 population crash, t-test, n.s. Colors correspond to seasons in panel A. **(F)** Seasonal graph clustering coefficients (average of all node specific clustering coefficients) in the 2 years pre and post removal of 20 RFID boxes, excluding seasons from the 2021 population crash, t-test, p < 0.01. Colors correspond to seasons in panel A. **(G)** Small world coefficients for all graphs in panel A by season one sample t-test by season comparing to a small world coefficient of 1, p < 0.01.

**Supplemental Figure 3.**
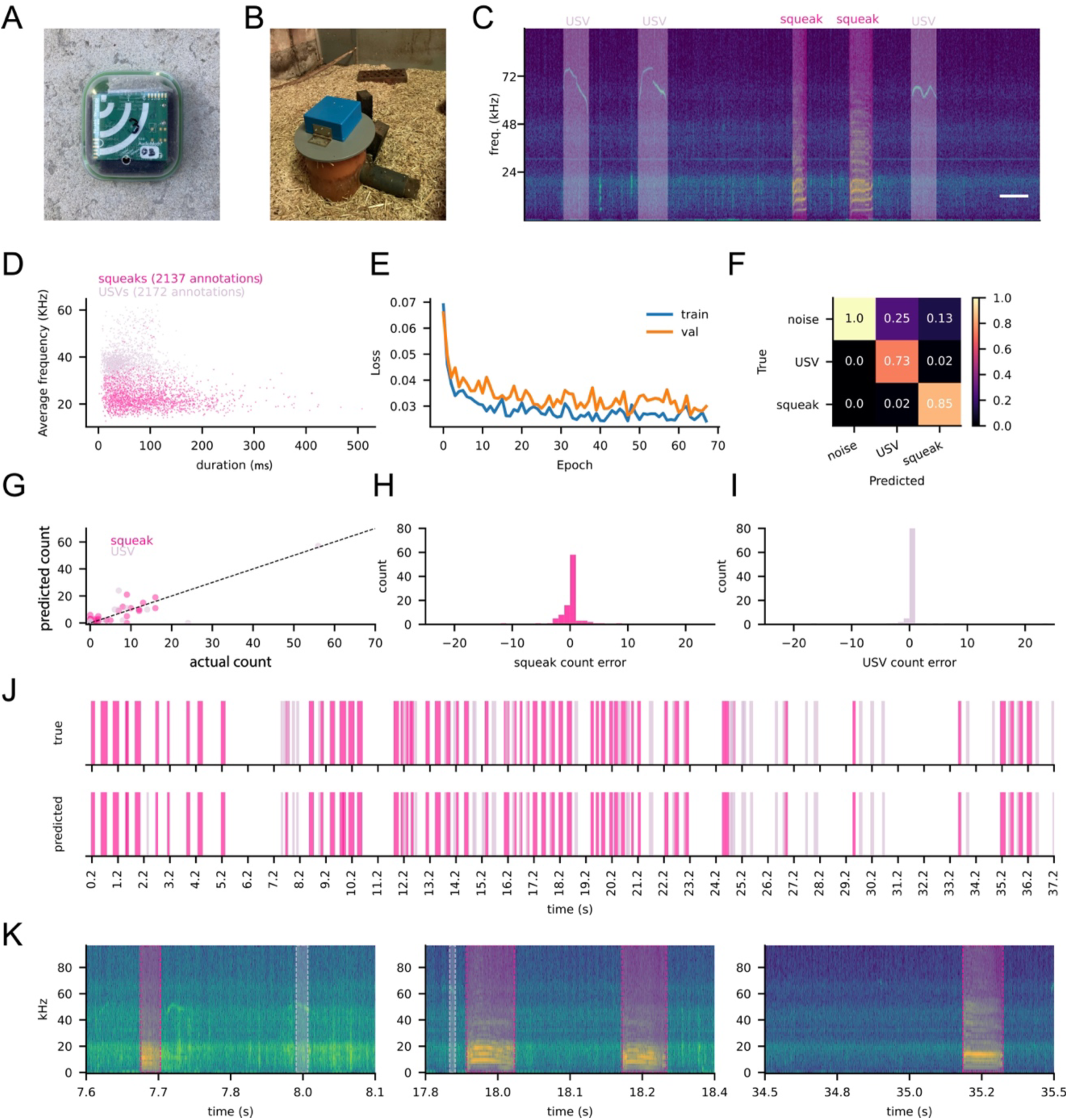
Evaluation of vocal segmentation model. (A) AudioMoth in custom case (B) AudioMoth recording from a box inside a protective container. (C) Example annotated spectrogram from an AudioMoth recording illustrating squeak and USV annotations. Scale bar: 100 ms (D) Duration and average frequency in kHz of all annotated USVs and squeaks (E) Training (train) and validation (val) loss for segmentation model from DAS (F) Confusion matrix for squeak and USV segments (G) Predicted vs actual vocalization counts for hand annotated recordings not part of the model training set (H) Squeak count error distribution from panel G (difference between actual squeak count per minute and predicted squeak count per minute) (I) USV count error distribution from panel G (difference between actual USV count per minute and predicted USV count per minute) (J) Example annotation (top) and prediction (bottom) from a recording not part of the training set (K) Example spectrograms from this recording showing example predictions.

